# DeepTetrad: high-throughput analysis of meiotic tetrads by deep learning in plants

**DOI:** 10.1101/677351

**Authors:** Eun-Cheon Lim, Jaeil Kim, Jihye Park, Eun-Jung Kim, Juhyun Kim, Hyun Seob Cho, Dohwan Byun, Ian R Henderson, Gregory P Copenhaver, Ildoo Hwang, Kyuha Choi

**Affiliations:** Department of Life Sciences, Pohang University of Science and Technology, Pohang, Gyeongbuk, Republic of Korea; Department of Plant Sciences, University of Cambridge, Cambridge CB2 3EA, UK; Department of Biology and the Integrative Program for Biological and Genome Sciences, University of North Carolina at Chapel Hill, Chapel Hill, North Carolina, USA

## Abstract

Meiotic crossovers facilitate chromosome segregation and create new combinations of alleles in gametes. Crossover frequency varies along chromosomes and crossover interference limits the coincidence of closely spaced crossovers. Crossovers can be measured by observing the inheritance of linked transgenes expressing different colors of fluorescent protein in *Arabidopsis* pollen tetrads. Here we establish DeepTetrad, a deep learning-based image recognition package for pollen tetrad analysis that enables high-throughput measurements of crossover frequency and interference in individual plants. DeepTetrad will accelerate genetic dissection of mechanisms that control meiotic recombination.

## Main

Meiosis consists of two consecutive nuclear divisions and produces four haploid gametes from a single diploid cell in sexually reproducing eukaryotes^1^. In *Arabidopsis* male meiosis, ∼200– 250 meiotic DNA double-strand breaks (DSBs) are induced in the genome by a DNA topoisomerase VI-like complex to initiate meiotic recombination^2–4^. Of these DSBs, only ∼8– 11 are repaired as crossovers (COs) using a homologous chromosome (homolog). Thus, male meiosis in the *Arabidopsis* genome, which comprises five chromosomes, results in an average of ∼1.8 crossovers between homologs. This low number suggests the existence of mechanisms that limit crossovers, a phenomenon that is observed in most eukaryotes^2^. Meiotic DSB and CO frequencies are controlled by genetic and epigenetic factors and are non-randomly distributed along chromosomes, with higher levels around gene promoters and terminators and lower levels across the centromeres^5–7^.

At least two pathways (Type I and Type II), contribute to CO formation^2,3^. The Type I pathway leads to interfering COs that prevent the coincident occurrence of closely spaced CO on the same pair of chromosomes^2,8,9^. In plants, interfering COs represent ∼80–85% of total COs and are dependent on the ZMM proteins (ZIP4, MSH4, MSH5, MER3, HEI10, SHOC1, PTD, MLH1, MLH3). The remaining ∼10–15% of COs are non-interfering and occur via the Type II pathway^10^. Non-interfering COs are resolved by the MUS81 endonuclease and are restricted by anti-recombination factors such as FANCM, RECQ4A, RECQ4B, and FIGL1^11–15^. Disruption of anti-recombination factors can increase the number of Type II COs in plants, which has the potential to create new combinations of desirable alleles that can improve crop varieties^13,16^. Therefore, high-throughput detection and understanding of CO frequency and interference have important implications for our understanding of the control of meiotic recombination as well as for breeding.

In *Arabidopsis*, CO frequency and interference can be measured by pollen tetrad analysis using Fluorescent-Tagged Lines (FTLs) in the *quartet1* (*qrt1*) background^17,18^. Mutation of the *QRT1* gene encoding a pectin methylestrase results in the four pollen products of male meiosis remaining attached to one another, allowing classical tetrad analysis. Each FTL has a transgenes that expresses eYFP (Y), dsRed (R) or eCFP (C) fluorescent proteins in mature pollen using the post-meiotic *LAT52* promoter. Genetic intervals bounded by transgenes expressing different colors (e.g. *I1bc, I1fg, I2ab, I2fg, I3bc, CEN3, I5ab*; Supplementary Fig. 1) can be created by crossing FTLs. Scoring the segregation of 2 or 3 linked markers enables CO frequency and interference to be measured. For example, plants that are hemizygous for the three markers (*YRC/+++*) that define the *I1bc* interval produce 12 pollen tetrad classes (A–L) depending on the number of COs between *YR* and *RC* (Supplementary Fig. 2). The relative segregation of any two markers can be used to place pollen tetrads into one of the three categories used for classic tetrad analysis: parental ditype (PD), tetratype (T) and non-parental ditype (NPD) (Supplementary Fig. 3). Tetrad analysis enables the calculation of map distances between pairs of markers, and measurement of CO interference between adjacent intervals.

Visual analysis of pollen tetrads is a powerful method for measuring genetic distance and crossover interference in *Arabidopsis*^12,14,15,17,18^. For example, manual analysis using fluorescence microscopy has been used to measure interference by comparing the map distances of two-color FTL intervals with and without COs in an adjacent interval^18^. However, manually scoring large numbers of tetrads is laborious and time consuming. Alternatively, *FTLs* in the *qrt1/+* or *QRT1* background can be analyzed by flow cytometry, which allows rapid measurement of CO frequency and interference of ∼10,000 single pollen grains per plant^19–21^. Unfortunately, the flow cytometric method is unable to detect double crossovers within single intervals, requires high purity pollen samples, and uses specialized equipment for three-color measurements. As an alternative, we have developed DeepTetrad (https://github.com/abysslover/deeptetrad), a deep learning-based image recognition package that enables quick, high-throughput, automated pollen tetrad analysis that can be used with existing FTL lines.

To develop DeepTetrad, we adapted the Mask Regional Convolutional Neural Network (Mask R-CNN), integrating a deep residual network (ResNet) backbone for image recognition to detect four-pollen tetrads with and without fluorescence^22,23^ (Fig. 1). First, DeepTetrad must precisely recognize pollen tetrads in bright-field pollen images, which include not only tetrads but also triads, dyads and monads (Fig. 1 a–c). DeepTetrad was assembled with two separate Mask R-CNN processes, using a ResNet-FPN backbone to generate masks of the bright-field pollen images (Fig. 1a)^23,24^. Then, DeepTetrad was trained to detect whole tetrad, triad, or dyad images via a Tetrad Segmentation Model with a ResNet depth of 101 layers. In parallel, we also trained DeepTetrad to detect every single pollen cell within tetrads, triads, dyads, and even monads via a Pollen Segmentation Model with a ResNet of depth 50 layers (Fig 1a). We used Keras and TensorFlow backends for training, with input bright-field images of pollen tetrads from *FTLs*^25,26^.

**Fig. 1.**
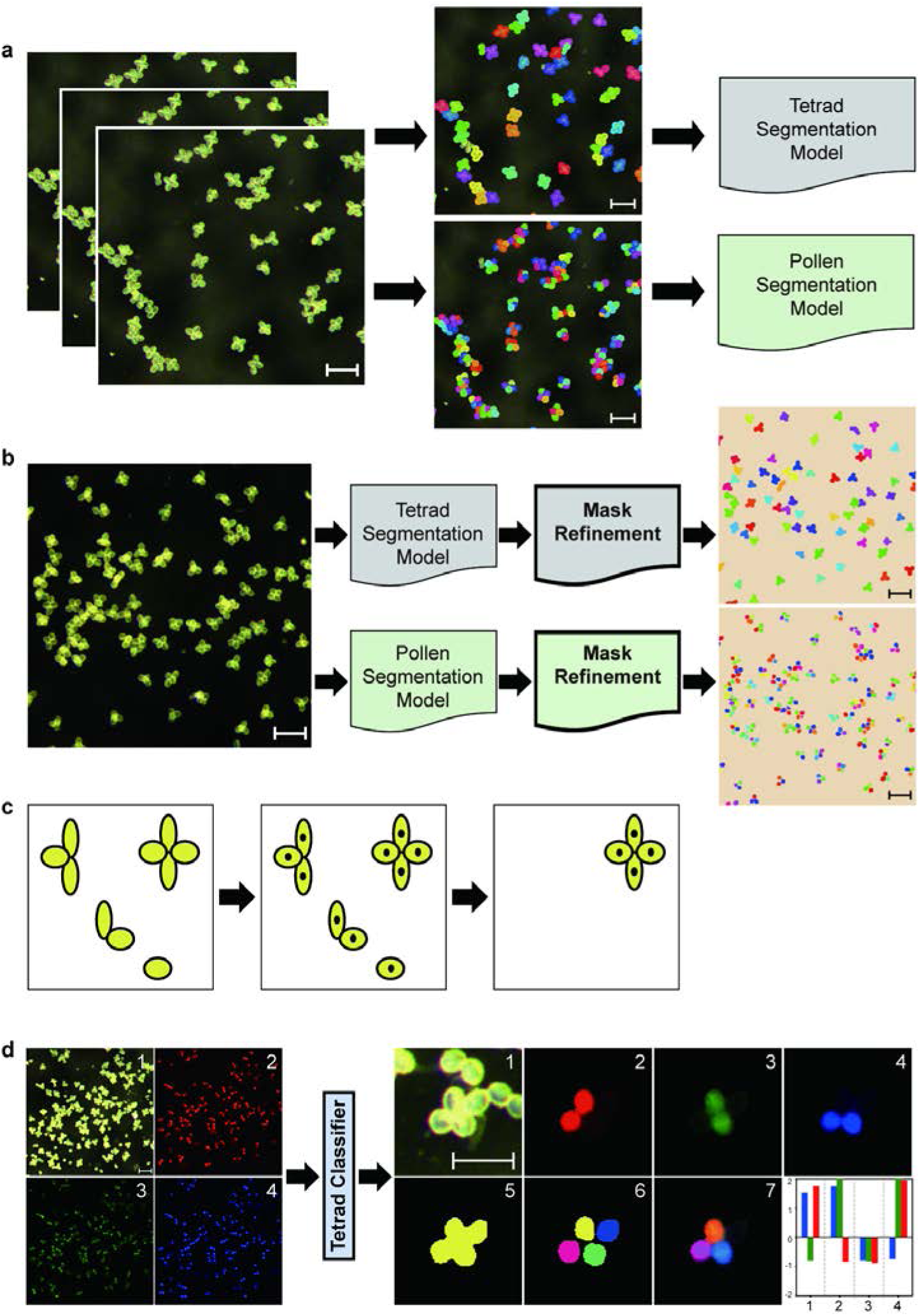
Establishment of DeepTetrad. **a,** Masking and training of tetrad-like and pollen images by DeepTetrad. Two separate DeepTetrad segmentation models make masks of tetrad-like and single pollens, respectively. **b,** Generation of masks from tetrad-like and single pollen images by DeepTetrad. **c,** Recognition and selection of measurable tetrad masks by DeepTetrad. Black dots represent the centroid assigned to each pollen mask in monads, dyads, triads, and tetrads. **d,** Tetrad classification by DeepTetrad. Bright-field (1), red (2), yellow (3), cyan (4)-filtered tetrad images, tetrad mask (5), single-pollen masks (6), three-color merged tetrads (7) and DeepTetrad output are displayed. In the bar graphs of DeepTetrad output, X axis labels indicate four pollens per tetrad and Y axis labels show the intensities of three-color fluorescence in the tetrad image. Scale bar = 0.1 mm (**a, b, d,** left), 0.05 mm (**d,** right). The colors in the tetrad mask images (**a,** middle panel, **b,** right panel, **d,** (6)) do not correspond to fluorescence colors.

When trained, DeepTetrad can produce masks of both tetrad-like (tetrads, triads, dyads) and single pollen-like objects from bright-field images of pollen tetrads (Fig. 1b). Next, DeepTetrad assigns a centroid to each pollen mask. Based on the position and distance between centroids of pollen masks within each tetrad-like mask, DeepTetrad recognizes measurable tetrads comprising four detectable pollen grains in the bright-field images (Fig. 1c). DeepTetrad‘s tetrad classifier then determines a tetrad type from a choice of 12 classes (A–L) for three-color assays (Supplementary Fig. 4), or from a choice of three types (PD, T, NPD) for two-color assays (Supplementary Fig. 5), according to the segregation pattern and intensity of fluorescence (yellow, red, cyan) in the four pollen masks per tetrad mask (Fig. 1d). CO frequency and interference can then be calculated using the frequency of tetrads in each class^18^. Because DeepTetrad is able to recognize single pollen grains and classify their fluorescence in tetrads, triads, dyads and monads, we developed the DeepMonad package by subclassing DeepTetrad. Like flow cytometry analysis, DeepMonad can measure crossover frequency and interference in FTLs by analyzing images of single fluorescent pollen grains in the *qrt1/+* and *QRT1* backgrounds^19^. In addition, DeepMonad can analyze single grains within tetrads, which allows comparison of genetic distances and interference calculated by DeepTetrad and DeepMonad (Fig. 2).

**Fig. 2.**
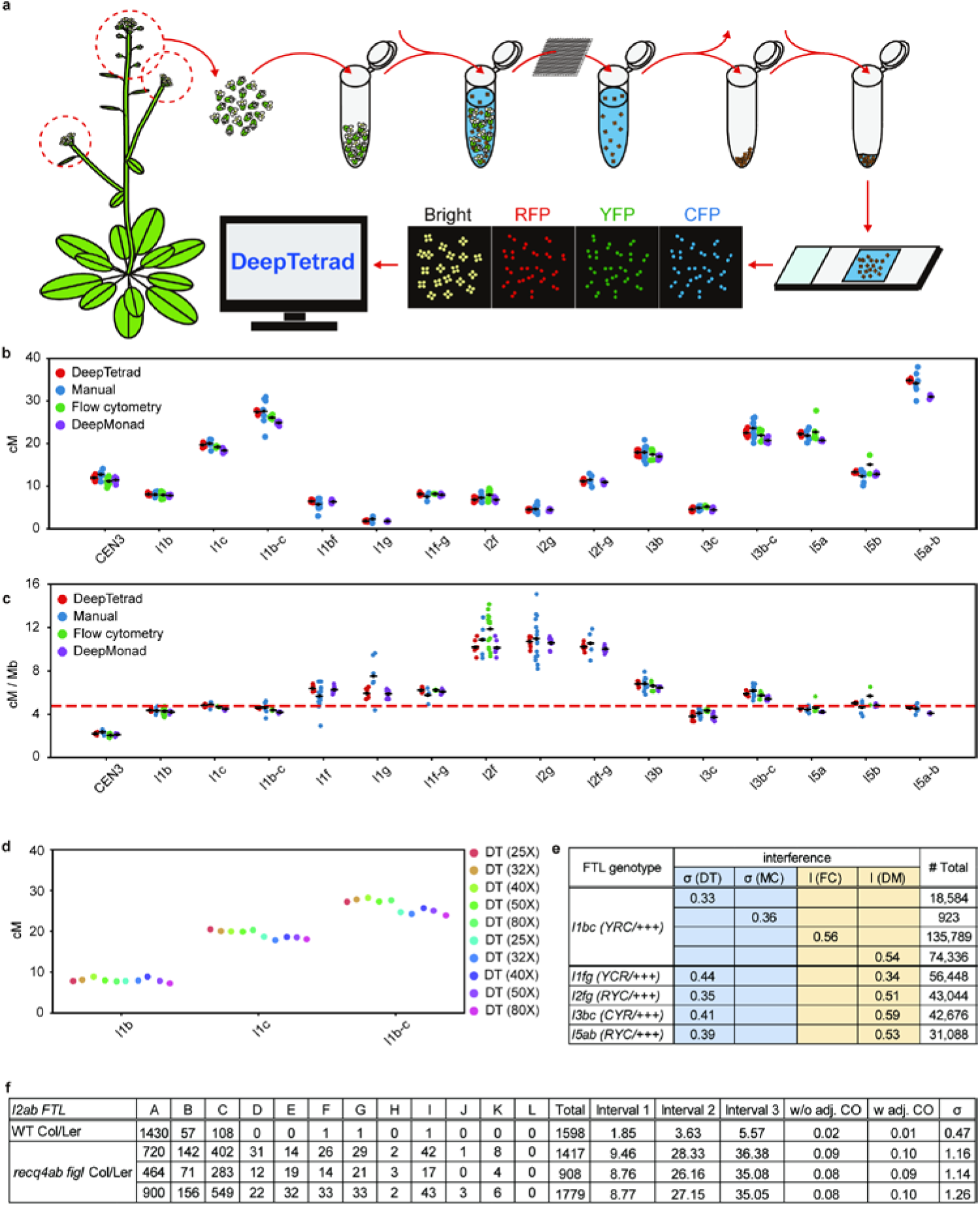
Measurements of crossover frequency and interference by DeepTetrad. **a,** A quick tetrad preparation method for high-throughput imaging of tetrads. The detail procedure was described in Methods. **b,** Plot showing measurement of CO frequencies (cM) in single intervals of FTLs. Genetic distances of single intervals were measured by DeepTetrad, manual counting, flow cytometry and DeepMonad. **c,** Plot showing measurement of CO frequencies (cM/Mb) in single intervals of FTLs. A horizontal red line indicates the male chromosome average crossover rate. **d,** Plot showing measurement of genetic distances in various sized tetrad images by DeepTetrad. Different magnifications were applied to the same tetrad samples for imaging (Supplementary Fig. 6). **e,** Measurement of CO interference by DeepTetrad. The CO interference ratio (σ= X*i1* without adjacent CO/ X*i1* with adjacent CO) was measured by DeepTetrad (DT) and manual counting (MC), highlighted in blue. Interference value (I=1-coefficenct of coincidence) in yellow, was calculated by flow cytometry (FC) and DeepMonad (DM). A value of 1 and 0 indicates no interference in σ and I, respectively. The values of interference in other FTLs were measured by DeepTetrad and DeepMonad. **f,** The CO interference ratio in *recq4a recq4b figl1* plants. DeepTetrad shows that *recq4a recq4b figl1* causes interference to be absent, increasing crossover frequency in FTL*-I2ab.*

Since DeepTetrad does not require specialized equipment (like flow cytometry), we developed a quick, simple method to prepare a large number of pollen tetrads for high-throughput imaging (Fig. 2a). This method allows extensive image sets of pollen tetrads (bright-field, red, yellow and cyan) to be obtained, which can then be analyzed quickly and simultaneously by DeepTetrad (Supplementary Table 1). We used this technique to measure genetic distances in two-color FTL intervals (*CEN3, I1b, I1c, I1b-c, I1f, I1g, I1f-g, I2f, I2g, I2f-g, I3b, I3c, I3b-c, I5a, I5b, I5a-b*) using DeepTetrad (Fig. 2b, Supplementary Fig. 1). The genetic distance values obtained this way were similar to those obtained using manual tetrad counting, flow cytometry and DeepMonad (Fig. 2b, Supplementary Table 2-4). Intriguingly, our DeepTetrad analysis showed higher crossover frequencies for long intervals (*I1b, I1c, I1b-c, I3b-c, I5a, I5b, I5a-b*) compared to DeepMonad single pollen analysis; this was because DeepTetrad, but not DeepMonad, detects double crossovers in long intervals (Fig. 2b, Supplementary Table 2-4). In addition, the *CEN3* interval which spans the centromere on chromosome 3 had a lower CO rate (2.21 cM/Mb) than the overall male chromosome average CO frequencies (4.77 cM/Mb), and intervals *I2f, I2g* and *I2f-g* which are close to the telomere had higher CO frequencies (10.19 cM/Mb, 10.71 cM/Mb and 10.23 cM/Mb, respectively) (Fig. 2c, Supplementary Fig. 1), consistent with prior observations^6,7,27^. DeepTetrad can also recognize tetrad images taken at different magnifications, and produces consistent CO frequency irrespective of scale (Fig. 2d, Supplementary Fig. 6).

To demonstrate DeepTetrad‘s utility for measuring CO interference we analyzed tetrad images from the three-color (*YRC/+++*) FTL interval-*I1bc* (Fig. 2e, Supplementary Fig. 1). Previously, interference had been measured in manually counted tetrads by calculating the interference ratio (defined here as σ) of the map distance of an interval (*i1*) in tetrads that have a CO in an adjacent interval (*i2*) with the map distance of the same interval (*i1*) in tetrads that lack a CO in *i2*^18^. Analysis of 18,584 tetrads with DeepTetrad resulted in a σ value of 0.33 for *I1bc* which is consistent with the σ value of 0.36 obtained by manually counting 923 tetrads (Fig. 2e, Supplementary Table 2-5). Flow cytometry has also been used to measure interference in fluorescent-tagged pollen monads by calculating the ratio of observed double COs to expected double COs (I=1-coefficient of coincidence, the ratio of DCO_obs_ to DCO_exp_)^19^. An analysis of 74,336 monads (converted from tetrad images) using this method with DeepTetrad resulted in an interference value of 0.54 for *I1bc* (*YRC/+++*) which was consistent with a value of 0.56 obtained from flow sorting 135,789 monads (Fig. 2e)^19^. To provide baseline values for future studies we also used DeepTetrad and DeepMonad to measured σ and I values in 4 other 3-color FTL intervals (*I1fg, I2fg, I3bc* and *I5ab*; (Fig. 2e, Supplementary Table 2 and 3). Previously, it was shown that frequency of Type II COs increases in *fancm* single mutants, as well as *recq4a recq4b figl1* triple mutants, leading to an absence of interference (I=0, σ =1)^12,14,15,19^. Using DeepTetrad, we found that the σ value of FTL*-I2ab* in *recq4a recq4b figl1* plants is 1, indicating no detectable interference, consistent with the prior observations (Fig. 2f, Supplementary Table 2 and 3). Taken together, our data demonstrate that DeepTetrad is a useful deep learning-based image recognition package for high-throughput measurements of both CO frequency and interference.

The FTL-based visual tetrad assay has been used extensively in studies of plant meiosis^12,17,18^. The application of flow cytometry to FTLs allowed rapid, high-throughput measurement of CO frequency and interference^19,21^. Here, we have extended the utility of FTLs further by developing DeepTetrad to enable quick, simple, and automated tetrad analysis. DeepTetrad will accelerate genetic analysis of meiotic recombination mechanisms as well as the influence of epigenetic and environmental effects.

## Methods

### DeepTetrad Network Architecture

DeepTetrad assembles two separate Mask Regional Convolutional Neural Network (Mask R-CNN) for the instance segmentation task^23^. They generate masks of pollen objects and tetrad objects, respectively, from input bright-field images. As backbone architectures, which are responsible for feature detection, we used deep residual networks (ResNet) of depths 50 and 101, with a feature pyramid network (FPN)^28^. We use the same terminology and definitions as those used in the Mask R-CNN article^23^ when describing the backbone; ResNet-50-FPN. Multi-task loss *L* is also defined in the same manner.

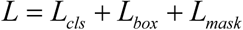

where *L*_*cls*_ is classification loss. *L*_*box*_ is loss of bounding box *v*=(*v*_*x*_, *v*_*y*_, *v*_*w*_, *v*_*h*_), which is a rectangle defined by coordinates of the upper-top vertex (*x, y*), and the dimension (width *w*, height *h*). *L*_*mask*_ is mask loss. The *i*-th mask of bounding box *v*, which is the core feature in DeepTetrad, is a grayscale image defined by the following logical predicate.

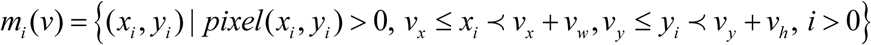

in which the *pixel*(*x, y*) function returns the pixel value of given coordinates in an image.

Mask segmentation is a multi-label classification. Hence *L*_*mask*_ must be calculated independently for each class in a single image. It is achieved by applying a sigmoid function to each pixel, from which the mean of binary cross entropy loss is calculated.

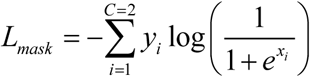

### Training

DeepTetrad is trained by nVidia TITAN X with 12 GB RAM using Keras^26^ and Tensorflow^25^ backends in the CUDA 10 platform. Transfer learning is performed with pre-trained weights on a Microsoft COCO dataset^29^. Input images are of fixed dimensions (1,920, 2,560). Zero-padding resizes each image to the exact dimension of (2,048, 2,560), which ensures that width and height are multiples of 512. The image is cropped at random positions with dimensions of (512, 512). For pollen objects, 919 training masks and 370 validation masks were used for data augmentation. For tetrad objects, 1,371 training masks and 617 validation masks were used. Masks were annotated using VGG Image Annotator (VIA)^30^.

Image augmentation with scaling, translation, rotation, and shearing operations is randomly triggered to each training session and validation mini-batch. During an epoch, a mini-batch of two images per Graphic Processing Unit (GPU) is fed to the backbone. Regions of interest (ROIs) or bounding boxes are sampled 128 times for the pollen model; 512 times for the tetrad model.

The pollen model, in which ResNet-50-FPN backbone is integrated, is trained by a single GPU for 10,000 iterations using the Stochastic Gradient Descent (SGD) optimizer with a learning rate of 0.001, momentum of 0.9, and weight decay of 0.0001. The tetrad model is trained with the same configuration, except that the backbone is replaced with ResNet-101-FPN and the number of iterations is increased to 20,000.

### Inference

Each backbone applies non-maximum suppression to 6,000 ROI candidates, in turn yielding 1,000 ROIs. The number of detected masks is limited by the maximum number of detecting instances *D*, which is set to 200. Hence, only *D*-detected ROIs with the highest scores are selected to create masks. If *D* is increased, the model may fail to infer masks because of memory limitation, or the inference may be seriously prolonged. The model might also overlook a considerable number of masks, which would be a critical problem.

As a solution, DeepTetrad tries to infer masks from cropped images rather than directly gathering them from a whole image. In an image of dimension (2048, 2560), there are at least 1,600–5,000 ground truth pollen masks, and at least 400–1,250 ground truth tetrad masks. As well as tetrad masks, monad, dyad, and triad masks will also be reported by the tetrad model. Thus, the number of ground truth masks in the whole image is much larger than *D* in both cases. A total of 63 cropped images of dimension (512, 512) are generated from the whole image, thereby left images of dimension (256, 512), or top images of dimension (512, 256) are intersected with one another. Complete inference for the whole image involves predicting masks from a mini-batch of 21 cropped images in three epochs.

### Mask refinement

All masks from the inference stage need to be refined. Masks that are produced at the edges of the cropped image are usually not overlapping in local coordinates, which represent the spatial location before translating to that in the whole image. However, they can become broken or overlapping when translated to global coordinates, which are the coordinates in the whole image. After removing broken or overlapping masks, contours and the number of tetrad and pollen masks can be more accurately determined. The procedure is as follows: i) all pixel coordinates of pollen masks, *M*_*p*_, and tetrad masks, *M*_*t*_ in each cropped image are moved to global coordinates by translation (affine transformation). ii) All pixel coordinates of tetrad masks are stored in a *k*-d tree, *T*_*t*_. Similarly, all pollen masks are kept in a *k*-d tree, *T*_*p*_. iii) All tetrad masks are queried against *T*_*t*_, and, similarly, all pollen masks are queried against *T*_*p*_ to collect refined tetrad masks, Ψt, and refined pollen masks, Ψp, which meet the predicate below:

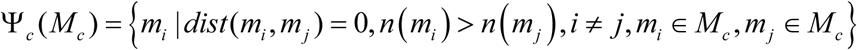

in which, *dist*(*x, y*) returns a Euclidean distance between *x* and *y, n*(*x*) returns the number of elements, and *c* is either *t* (tetrad) or *p* (pollen).

The measurable tetrad masks and pollen masks, Ω, are defined as follows:

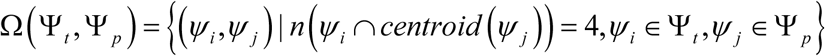

where *centroid*(*x*) yields the median of given mask coordinates.

Centroids are calculated to associate a tetrad mask Ψ_t_, with the refined pollen masks Ψ_p_. Each centroid of a pollen mask, Ψ_p_, is queried against *T*_*t*_, then Ψ_p_ is associated with a tetrad mask, Ψ_t_ if the Euclidean distance between them is 0. When the number of Ψ_p_ associated with Ψt is four, they are deemed to be measurable.

### Tetrad classification

A tetrad mask can be classified into a representative type of crossover event according to the fluorescence intensity values of associated pollen masks. The signal intensity, *S*_*c*_, is defined as the mean of pixel values in each fluorescence channel of measurable pollen masks, in which *c* can be a fluorescence channel of red (R), yellow (Y), or cyan (C). We assume *c*={R, Y} for two-channel images, and *c*={R, Y, C} for three-channel images. For measurable tetrad masks, *S*_*c*_ of four pollen masks can be calculated, yet those which have undergone silencing in fluorescent protein expression should be ignored. If silencing occurs, the difference between *S*_*c*_ of the second-highest and the third-highest would be smaller than a certain threshold, Θ. Second-highest *S*_*c*_ represents presence or on-state of fluorescence proteins, whereas the third-highest implies absence or off-state of the proteins. Z-scores are calculated for four *S*_*c*_ values before finding Θ. We explored a parameter space of Θ up to two decimal places using an adaptive grid search method, then Θ was set to 0.40, meaning that *Sc* differences between on-state and off-state must be bigger than but not equal to 0.40 in a normal distribution.

We determine if fluorescent proteins in a pollen grain are expressed by comparing individual *S*_*c*_ with the median of all four *S*_*c*_ values. With per-channel expressions, tetrad masks are classified as one of classical three tetrad types: parental ditype (PD), tetra type (T), and non-parental ditype (NPD), with two-color FTL intervals or 12 classes (A to L) with the three-color counterparts (Supplementary Fig. 2 and 3). In three-color FTL intervals that have two intervals (*i1* and *i2*) with four chromatids (1-4), tetrad classes are non-crossover (A), single crossover interval 1 (B; SCO-*i1*), single crossover interval 2 (C; SCO-*i2*), two strand double crossover (D; 2stDCO), three strand double crossover a (E; 3st DCOa), three strand double crossover b (F; 3st DCOb), four strand double crossover (G; 4st DCO), non-parental ditype interval 1, non-crossover interval 2 (H; NPD-*i1* NCO-*i2*), non-crossover interval 1, non-parental ditype interval 2 (I; NCO-*i1* NPD-*i2*), non-parental ditype interval 1, single crossover interval 2 (J; NPD-*i1* SCO-*i2*), single crossover interval 1, non-parental ditype interval 2 (K; SCO-*i1* NPD-*i2*) and non-parental ditype interval 1, non-parental ditype interval 2 (L; NPD-*i1* NPD-*i2*)^18^.

### Calculation of interference

With two-color FTL intervals, we calculate crossover frequency following Perkin‘s equation:

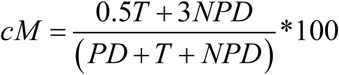

With three-color FTL intervals, we can calculate the interference ratio σ, which is the ratio of the map distance with adjacent crossover *χ*_*γ*_ to the map distance without adjacent crossover *X*_*δ*_

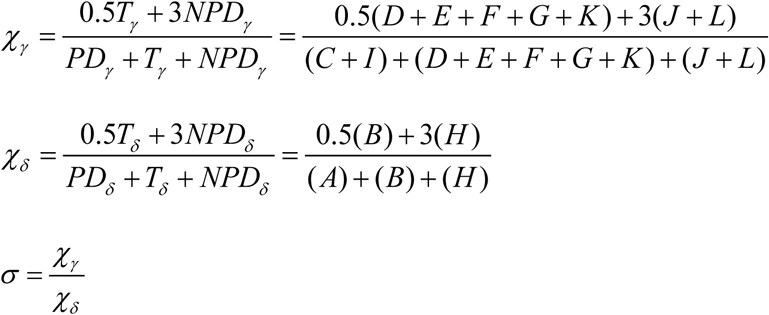

in which PD means the number of parental ditypes or no crossover event. T is the number of tetratype or single crossover events. NPD is the number of non-parental ditype or double crossover events. A-L letters represent 12 tetrad classes from three-color FTLs (Supplementary Fig. 2)^18^.

In the above equations, we assumed that two adjacent genetic intervals of *i1* and *i2* are defined by three separate fluorescent protein transgenes of red, yellow, and cyan in sequential order. The γ represents that at least a single crossover event occurred at *i2*, meanwhile_*δ*_denotes that no crossover events were found at *i2*. We highlight that the number of crossover event is counted at *i1*. Tγ is the tetratype tetrads for *i1* that have a CO in *i2*, and T_*δ*_ is the tetratype tetrads for *i1* that do not have a CO in *i2*. DeepTetrad maps the order of input color images (red-yellow-cyan) to the physical order of fluorescent protein transgenes of FTLs for calculating the interference ratio as well as genetic distance.

### Pollen tetrad preparation

FTL plants were grown at 20°C under long-day condition (16 h light/8 h dark). Twenty open flowers of a primary shoot from 30-day old FTL plants were collected in a 1.5-ml tube, and 1 ml of pollen tetrad preparation solution (17% sucrose, 2 mM CaCl_2_, 1.625 mM boric acid, 0.1% Triton-X-100, pH 7.5) was added before incubating for 5 min at room temperature, with gentle rotation. Flowers and the solution were mixed by inverting the tube several times. The solution of pollen tetrads was pipetted and filtered into a new 1.5-ml tube through an 80-μm nylon mesh (30 × 30 mm). The filtered solution was centrifuged at 500*g* for 3 min to make a yellow pellet. The supernatant was removed and discarded by pipetting or vacuum aspiration. Four μl of pollen tetrad preparation solution was added to the yellow pellet. After pipetting gently five times, the 4-μl suspension of pollen tetrads was loaded on a glass microscope slide and covered with a small cover glass (9 × 9 mm). This resulted in ∼2,500 tetrads for imaging.

### Microscopy and imaging

A set of four photographs for each pollen tetrad was taken using a Leica M165 FC dissecting stereomicroscope with bright-field, RFP, YFP and CFP filters (ArticleNo 10450224, 10447410, 10447409, respectively) in sequential order. Twelve image sets per cover glass were obtained from ∼20 flowers when a magnification of 50x was used to image tetrads. Information about the gain, gamma, saturation and exposure for each FTL for high quality imaging is available (Supplementary Table 6).

## Acknowledgements

We thank Raphael Mercier (MPI, Germany) for providing *recq4 figl1* mutant seeds. We thank our colleagues for giving critical comments. This work is funded by the Suh Kyungbae Foundation, Republic of Korea, Next-Generation BioGreen 21 Program (PJ01337001), Rural Development Administration, Republic of Korea, and Basic Science Research Program through the National Research Foundation of Korea (NRF) funded by the Ministry of Education (2017R1D1AB03028374). GPC is supported by a US National Science Foundation grant (IOS-1844264).

---

**Supplementary Fig. 1.**
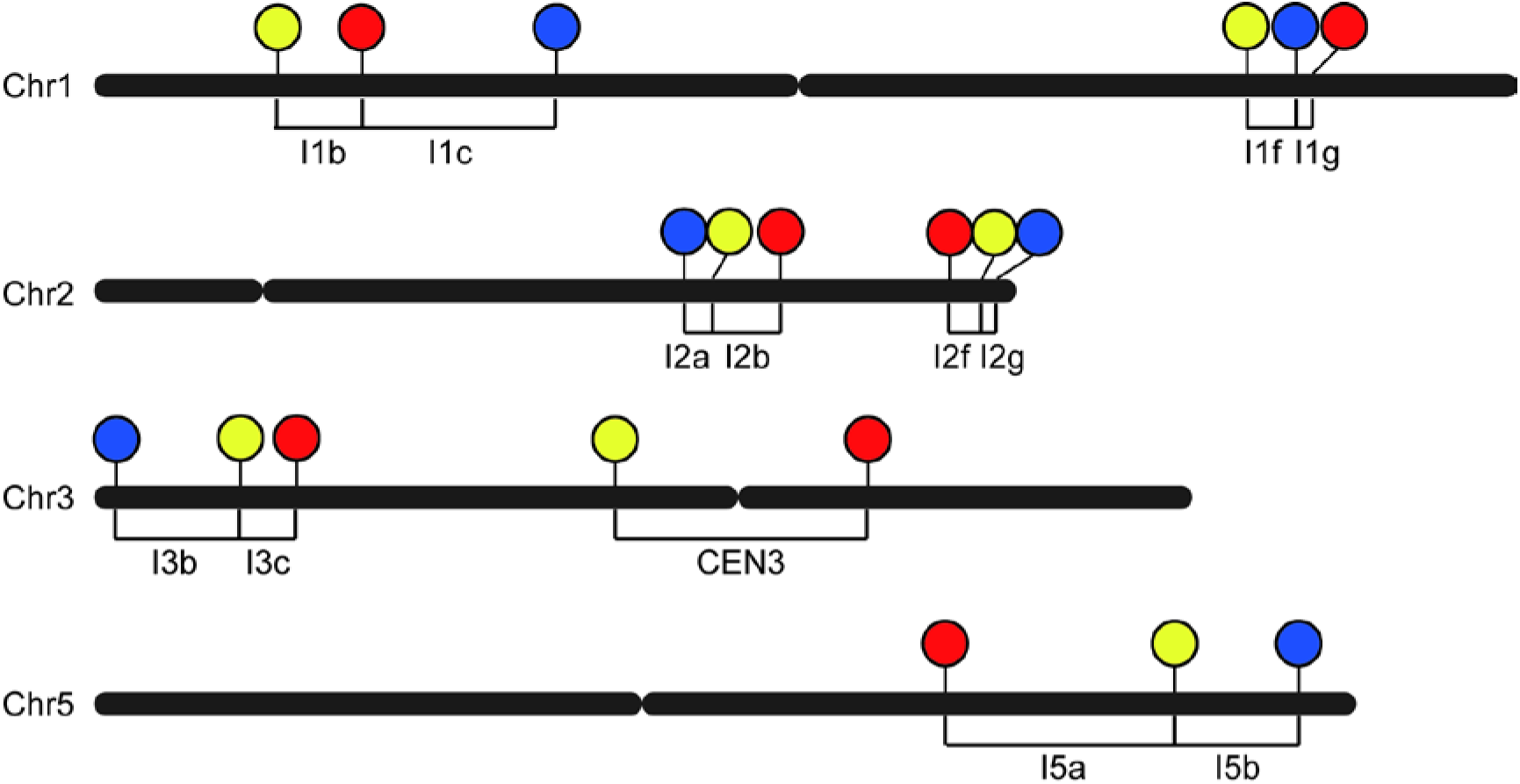
T-DNA locations of pollen FTLs (*I1bc, I1fg, I2ab, I2fg, I3bc, CEN3, I5ab*) on the *Arabidopsis thaliana* genome. T-DNA positions of FTLs originally generated in *qrt1* Col-0 plants are displayed on the *Arabidopsis* genome. Each FTL of homozygous genotype for fluorescence T-DNAs was crossed to *qrt1*, and pollen tetrads of F_1_ plants were used to measure CO frequency and interference by DeepTetrad. Chr = chromosome.

**Supplementary Fig. 2.**
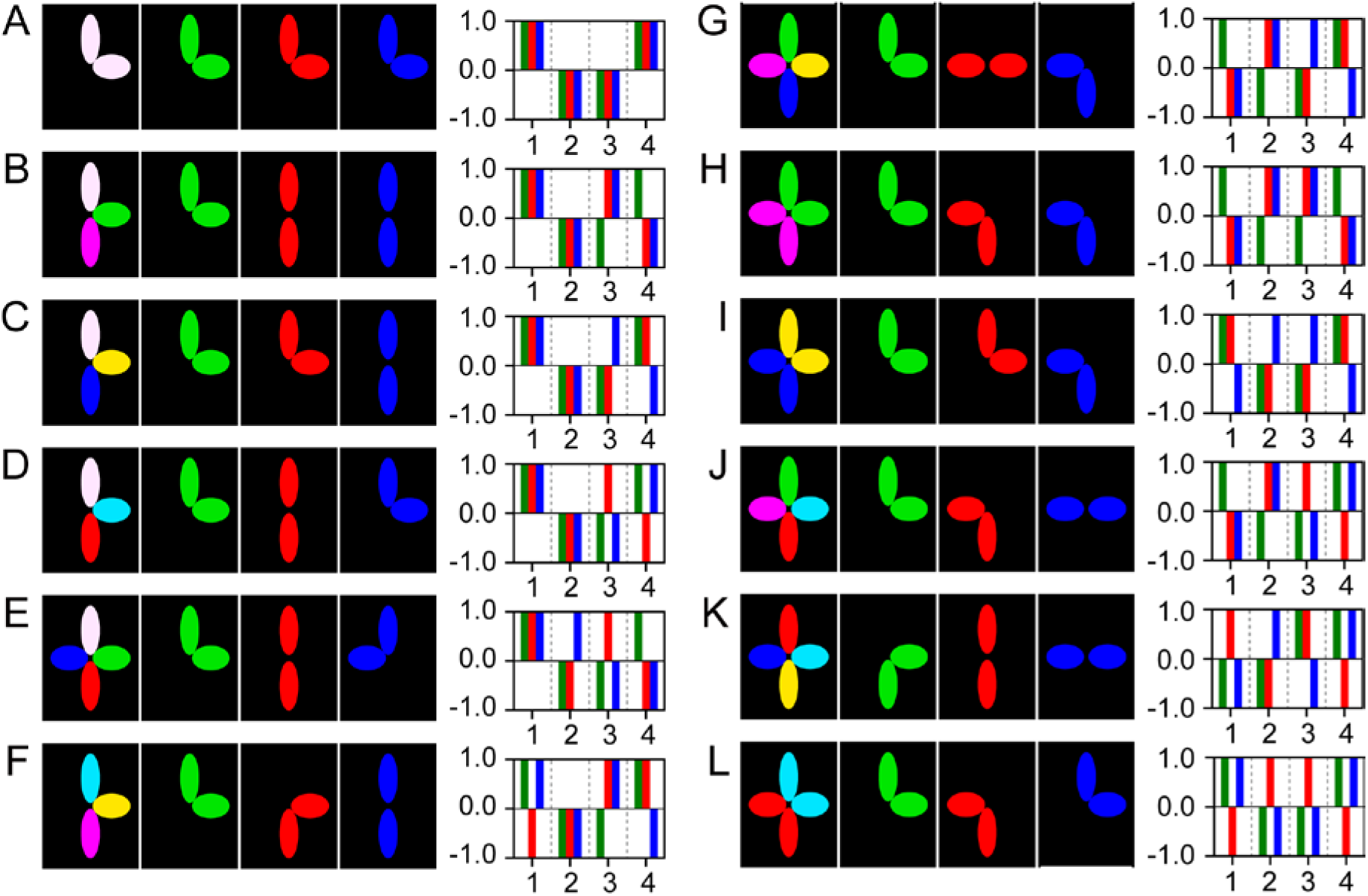
Diagram of 12 tetrad classes (A–L) generated by three-color tetrad assay and DeepTetrad outputs. Twelve tetrad classes (**A**-**L**) are generated from three-color tetrad assays^18^. According to the position and segregation of three T-DNAs expressing eYFP, dsRed and eCFP, non-crossover(**A**) and recombination tetrad classes (**B**-**L**) are determined in FTLs (*YRC/+++*). For each tetrad type, merged, red, yellow and cyan images are displayed. Bar graphs indicate the output of DeepTetrad classification. In the bar graphs, X axis labels indicate four pollens (1, 2, 3, 4) per tetrad and Y axis labels show the intensities of three-color fluorescence in the tetrad images.

**Supplementary Fig. 3.**
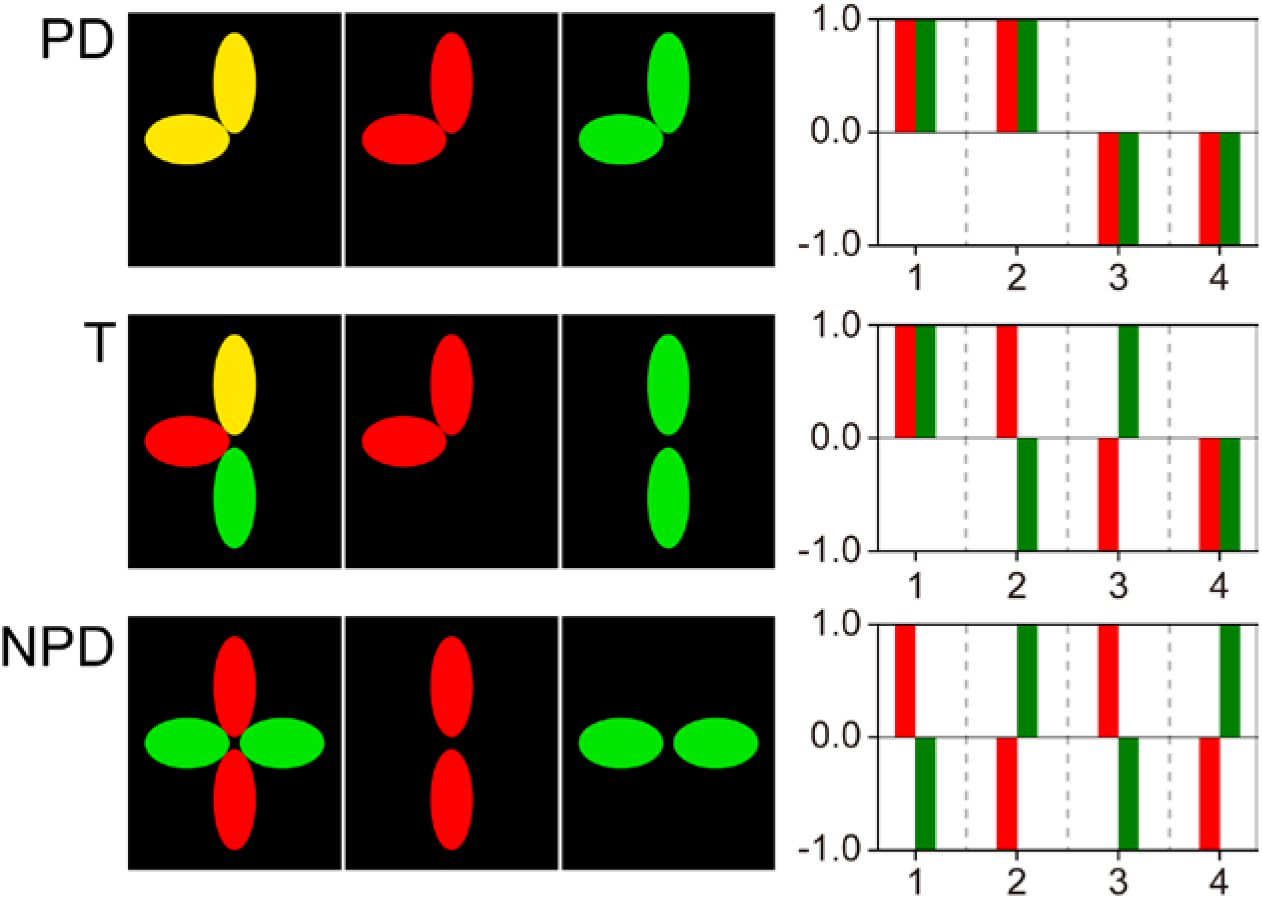
Diagram for three tetrad classes (PD, T, NPD) of 2-color assay and DeepTetrad output. Tetrad classes of **PD** (parental ditype), **T** (tetra type), and **NPD** (non-parental ditype) are generated from two-color tetrad assays of FTLs (*RY/++*). In the bar graphs, X axis labels indicate four pollens per tetrad and Y axis labels show the intensities of two-color fluorescence in the tetrad images.

**Supplementary Fig. 4.**
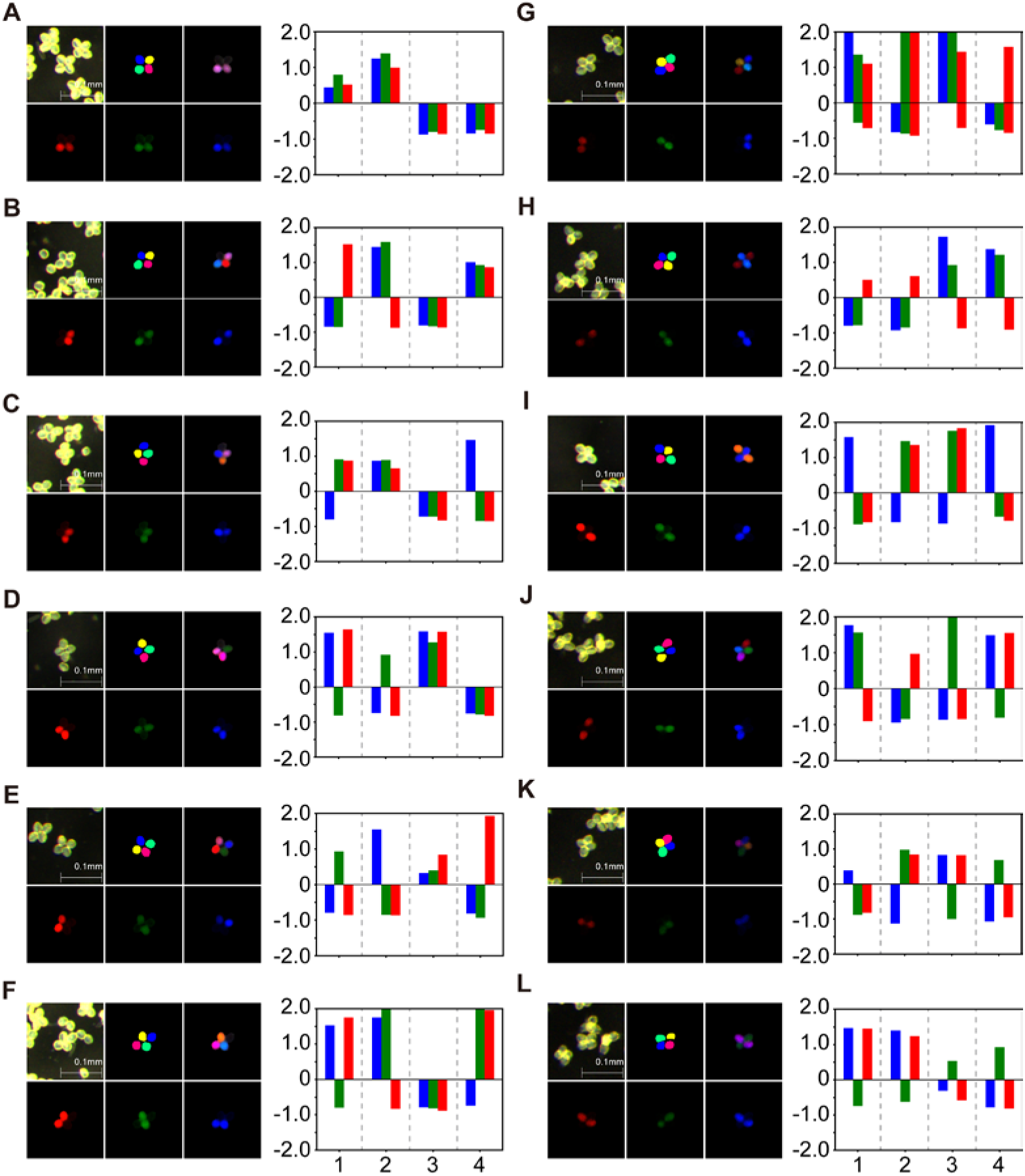
Tetrad images and DeepTetrad output in three-color assay of FTL plants. DeepTetrad recognizes measurable tetrads from a large number of tetrads, and classifies the tetrads as **A** (non-crossover, NCO) or **B**–**L** (recombinant tetrad classes with one or two crossovers in interval 1 and 2 (*i1* and *i2*)) of FTL (*RYC/++*). Tetrad classes are **B**, single crossover interval 1 SCO-*i1,* **C**, SCO-*i2,* **D**, two strand double crossover 2st DCO, **E**, 3st DCOa, **F**, 3st DCOb, **G**, 4st DCO, **H**, NPD-*i1* NCO-*i2*, **I**, NCO-*i1* NPD-*i2*, **J**, NPD-*i1* SCO-*i2*, **K**, SCO-*i1* NPD-*i2*, and **L**, NPD-*i1* NPD-*i2*^18^. Each panel (A-L) shows bright-field (upper left), single-pollen mask (upper middle), merged-fluorescent (upper right), and single-color fluorescent (lower three) images. The colors in the tetrad mask images do not correspond to fluorescence colors. In the bar graphs, X axis labels indicate four pollens per tetrad and Y axis labels show the intensities of three-color fluorescence in the tetrad images. Scale bar = 0.1 mm.

**Supplementary Fig. 5.**
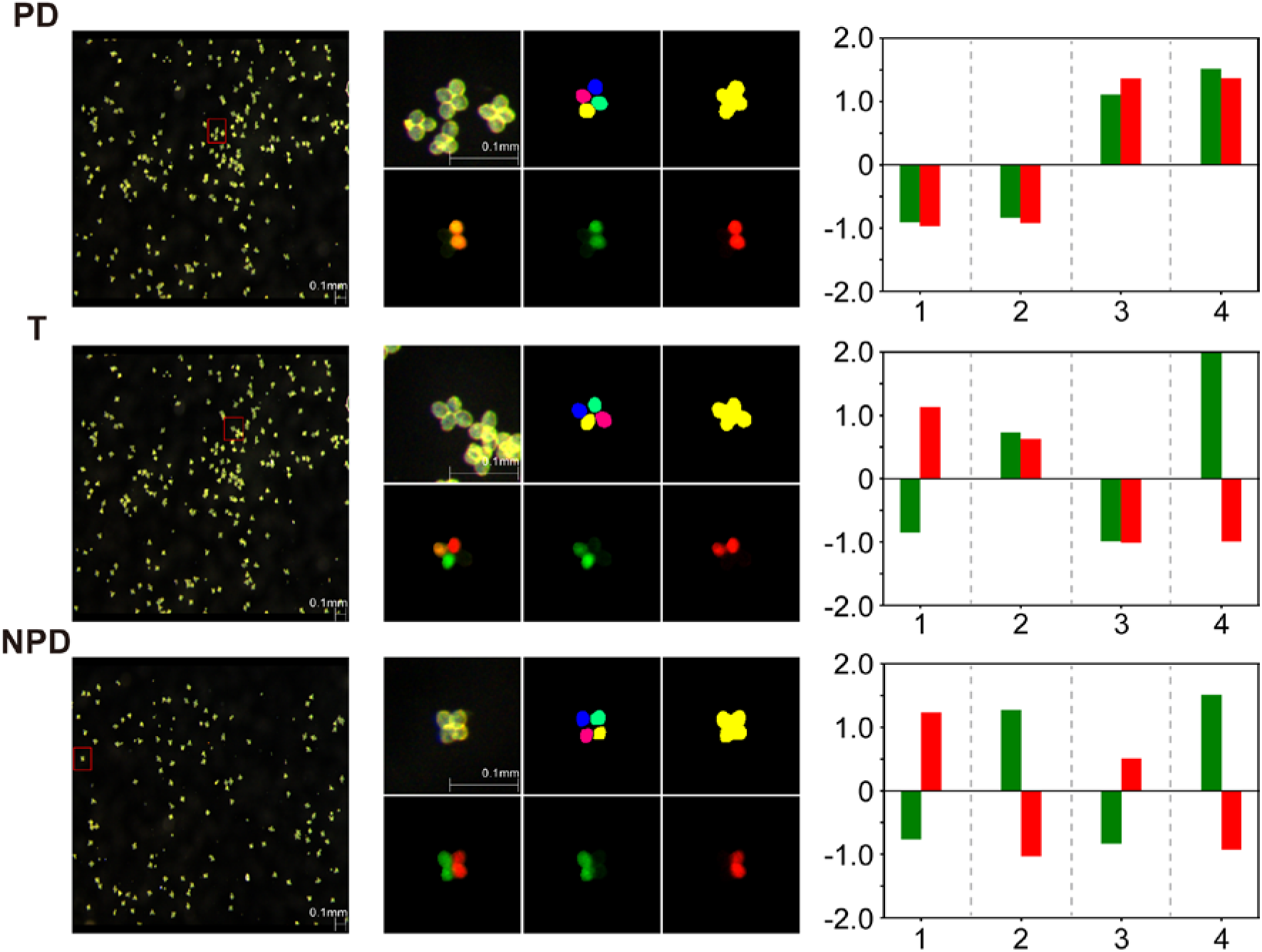
Tetrad images and DeepTetrad output in two-color assay of FTL*-CEN3 (YR/++)* plants. DeepTetrad recognizes measurable tetrads (left panels) and classifies them as **PD** (parental ditype), **T** (tetra type) and **NPD** (non-parental ditype) of tetrad types according to segregation of fluorescence in FTL*-CEN3* (*YR/++*) (middle panels). Each panel in the middle shows bright-field (upper left), single-pollen mask (upper middle), tetrad mask (upper right), merged-fluorescent (lower left), yellow (lower middle) and red (lower right) fluorescent images. The colors in the tetrad mask images do not correspond to fluorescence colors. In the bar graphs, X axis labels indicate four pollens per tetrad and Y axis labels show the intensities of two-color fluorescence in the tetrad images. Scale bar = 0.1 mm.

**Supplementary Fig. 6.**
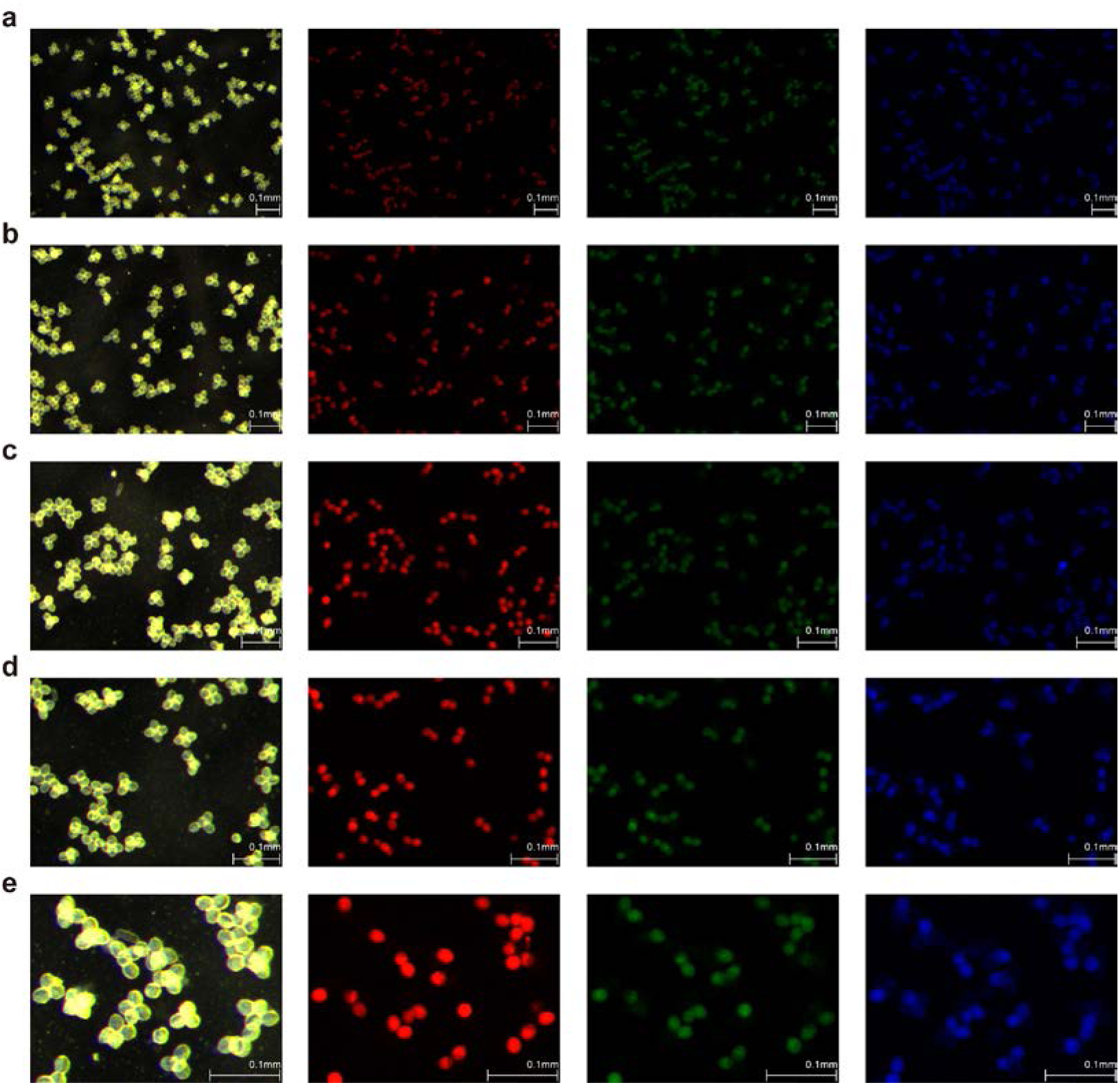
Different sized tetrad images. Tetrad images were taken using an epifluorescence microscope at different magnifications (**a**– **e**) (25x, 32x, 40x, 50x, 80x) under bright-field, RFP, YPF and CFP filters. Scale bar = 0.1 mm.

**Supplementary Table 1.** Comparison of crossover measurement methods. DeepTetrad involves the preparation of a large number of tetrads using a quick and simple method, and increases the speed of tetrad analysis to obtain data from many individual plants by analyzing all tetrad images simultaneously. Abbreviations: CO, crossover; DCO, double crossovers.

|  | **Tetrad analysis**<br>**(manual counting)** | **Flow cytometry** | **Tetrad analysis**<br>**(DeepTetrad)** |
| --- | --- | --- | --- |
| Equipment requirements | Fluorescence microscope, graphics software | Fluorescence microscope, flow cytometer | Fluorescence microscope, DeepTetrad package |
| Tetrad preparation method | Simple | Multiple steps (50-ml tube, flow cytometric tube) | Quick and simple (1.5-ml tube) |
| Time to prepare tetrads from 10 individual plants | 2 h 30 min (1 h 40 min/sampling, 50 min/imaging) | 1 h (60 min/sampling) | 1 h (10 min/sampling, 50 min/imaging) |
| Time of data analysis for two adjacent intervals in 10 individual plants (~1,000 tetrads/plant) | 30 h | 5 h (30 min/plant, 10 plants) | 2 h 30 min (16 min/plant, 10 plants) |
| Imaging requirement | Yes | No | Yes |
| Data analysis | Individually | Individually | Simultaneously |
| Gene conversion measurement | Yes | No | Yes |
| CO interference measurement | Yes | Yes | Yes |
| Single-interval DCO measurement | Yes | No | Yes |
| Differentiating 2-strand, 3-strand and 4-strand DCOs from one another | Yes | No | Yes |
| *qrt1* mutant background | Yes | No | Yes |

**Supplementary Table 2.** Measurement of crossover frequency of FTL intervals by DeepTetrad, manual counting, flow cytometry, and DeepMonad. In FTLs with T-DNAs expressing eYFP (Y), dsRed (R) and eCFP (C), the physical distances and genetic distances are shown^18^. Genetic distances were measured by DeepTetrad, manual counting, flow cytometry^6,19–21,31^, and DeepMonad. The results of statistical analyses (mean of cM, 95% confidence interval, *P*-value) on the genetic distances measured by different methods are shown. The *P*-values of significantly different crossover frequency are marked by asterisks. A *P*-value is not calculated when assumptions for the statistical *t*-test are violated. Abbreviations: DT, DeepTetrad; MC, manual counting; FC, flow cytometry; DM, DeepMonad; FTL, fluorescent tagged line.

| FTL genotype | T-DNA 1 | T-DNA 2 | T-DNA 3 | Mb | cM (DT) | cM (MC) | cM (FC) | cM (DM) | *p*-value (DT-MC) | *p*-value (DT-DM) | Total tetrads (DT) |
| --- | --- | --- | --- | --- | --- | --- | --- | --- | --- | --- | --- |
| *l1bc* (YRC/+++) | 3,905,441 | 5,755,618 | 9,850,022 |  |  |  |  |  |  |  | 18,584 |
| *l1b* (YR/++) | 3,905,441 | 5,755,618 |  | 1.85 | 8.09 ± 0.17 | 8.02 ± 0.59 | 7.93 ± 0.18 | 7.79 ± 0.20 | 7.68E-01 | 1.54E-02* |  |
| *l1c* (RC/++) |  | 5,755,618 | 9,850,022 | 4.09 | 19.69 ± 0.37 | 20.00 ± 0.83 | 19.20 ± 0.21 | 18.38 ± 0.32 | 3.97E-01 | 2.02E-05* |  |
| *l1b-c* (YC/++) | 3,905,441 |  | 9,850,022 | 5.94 | 27.43 ± 0.36 | 27.59 ± 3.10 | 26.12 ± 0.42 | 24.88 ± 0.32 | 9.07E-01 | 5.29E-09* |  |
| *l1fg* (YCR/+++) | 24,645,163 | 25,652,977 | 25,956,590 |  |  |  |  |  |  |  | 14,112 |
| *l1f* (YC/++) | 24,645,163 | 25,652,977 |  | 1.01 | 6.44 ± 0.21 | 5.74 ± 0.5 |  | 6.34 ± 0.26 | 5.61E-02 | 4.78E-01 |  |
| *l1g* (CR/++) |  | 25,652,977 | 25,956,590 | 0.30 | 1.78 ± 0.11 | 2.26 ± 0.61 |  | 1.77 ± 0.10 | 1.02E-01 | 8.19E-01 |  |
| *l1f-g* (YR/++) | 24,645,163 |  | 25,956,590 | 1.31 | 8.14 ± 0.18 | 7.54 ± 2.5 | 8.16 ± 0.07 | 7.95 ± 0.18 | 1.16E-01 | 1.05E-01 |  |
| *l2fg* (RYC/+++) | 18,286,716 | 18,957,093 | 19,373,634 |  |  |  |  |  |  |  | 10,761 |
| *l2f* (RY/++) | 18,286,716 | 18,957,093 |  | 0.67 | 6.83 ± 0.33 | 7.29 ± 1.3 | 7.95 ± 0.58 | 6.79 ± 0.34 | 3.94E-01 | 8.63E-01 |  |
| *l2g* (YC/++) |  | 18,957,093 | 19,373,634 | 0.42 | 4.50 ± 0.17 | 4.61 ± 0.46 |  | 4.45 ± 0.18 | 6.31E-01 | 6.08E-01 |  |
| *l2f-g* (RC/++) | 18,286,716 |  | 19,373,634 | 1.09 | 11.15 ± 0.28 | 11.5 ± 2.31 |  | 10.93 ± 0.26 | 6.67E-01 | 1.93E-01 |  |
| *l3bc* (CYR/+++) | 498,916 | 3,126,994 | 4,319,513 |  |  |  |  |  |  |  | 10,669 |
| *l3b* (CY/++) | 498,916 | 3,126,994 |  | 2.63 | 17.91 ± 0.54 | 17.94 ± 0.71 | 17.43 ± 1.14 | 16.95 ± 0.35 | 9.58E-01 | NA |  |
| *l3c* (YR/++) |  | 3,126,994 | 4,319,513 | 1.19 | 4.53 ± 0.35 | 4.88 ± 0.28 | 5.17 ± 0.14 | 4.44 ± 0.31 | 8.17E-02 | 6.58E-01 |  |
| *l3b-c* (CR/++) | 498,916 |  | 4,319,513 | 3.82 | 22.51 ± 0.64 | 23.61 ± 1.46 | 21.89 ± 1.22 | 20.72 ± 0.53 | 1.44E-01 | 1.65E-04* |  |
| *l5ab* (RYC/+++) | 18,164,269 | 23,080,567 | 25,731,311 |  |  |  |  |  |  |  | 7,772 |
| *l5a* (RY/++) | 18,164,269 | 23,080,567 |  | 4.92 | 22.24 ± 0.49 | 21.86 ± 1.36 | 22.70 ± 1.77 | 20.73 ± 0.27 | 5.32E-01 | 7.28E-05* |  |
| *l5b* (YC/++) |  | 23,080,567 | 25,731,311 | 2.65 | 13.31 ± 0.31 | 12.36 ± 1.28 |  | 12.81 ± 0.42 | 1.62E-01 | 2.68E-02* |  |
| *l5a-b* (RC/++) | 18,164,269 |  | 25,731,311 | 7.57 | 34.91 ± 0.38 | 34.21 ± 2.41 |  | 31.03 ± 0.50 | 5.05E-01 | 1.35E-07* |  |
| *CEN3* (RY/++) | 11,115,724 | 16,520,560 |  | 5.40 | 11.96 ± 0.64 | 12.72 ± 0.86 | 11.17 ± 0.19 | 11.45 ± 0.70 | 1.16E-01 | 1.95E-01 | 7,657 |
| *l2ab recq4ab figl* (CYR/+++) | 12,640,092 | 13,226,013 | 14,675,407 |  |  |  |  |  |  |  | 4,271 |
| *l2a recq4ab figl* (CY/++) | 12,640,092 | 13,226,013 |  | 0.59 | 8.86 ± 1.38 |  |  | 8.35 ± 1.56 | NA | NA |  |
| *l2b recq4ab figl* (YR/++) |  | 13,226,013 | 14,675,407 | 1.45 | 27.47 ± 1.88 |  |  | 21.42 ± 0.48 | NA | NA |  |
| *l2a-b recq4ab figl* (CR/++) | 12,640,092 |  | 14,675,407 | 2.04 | 35.26 ± 2.56 |  |  | 25.99 ± 0.83 | NA | NA |  |

**Supplementary Table 3.**
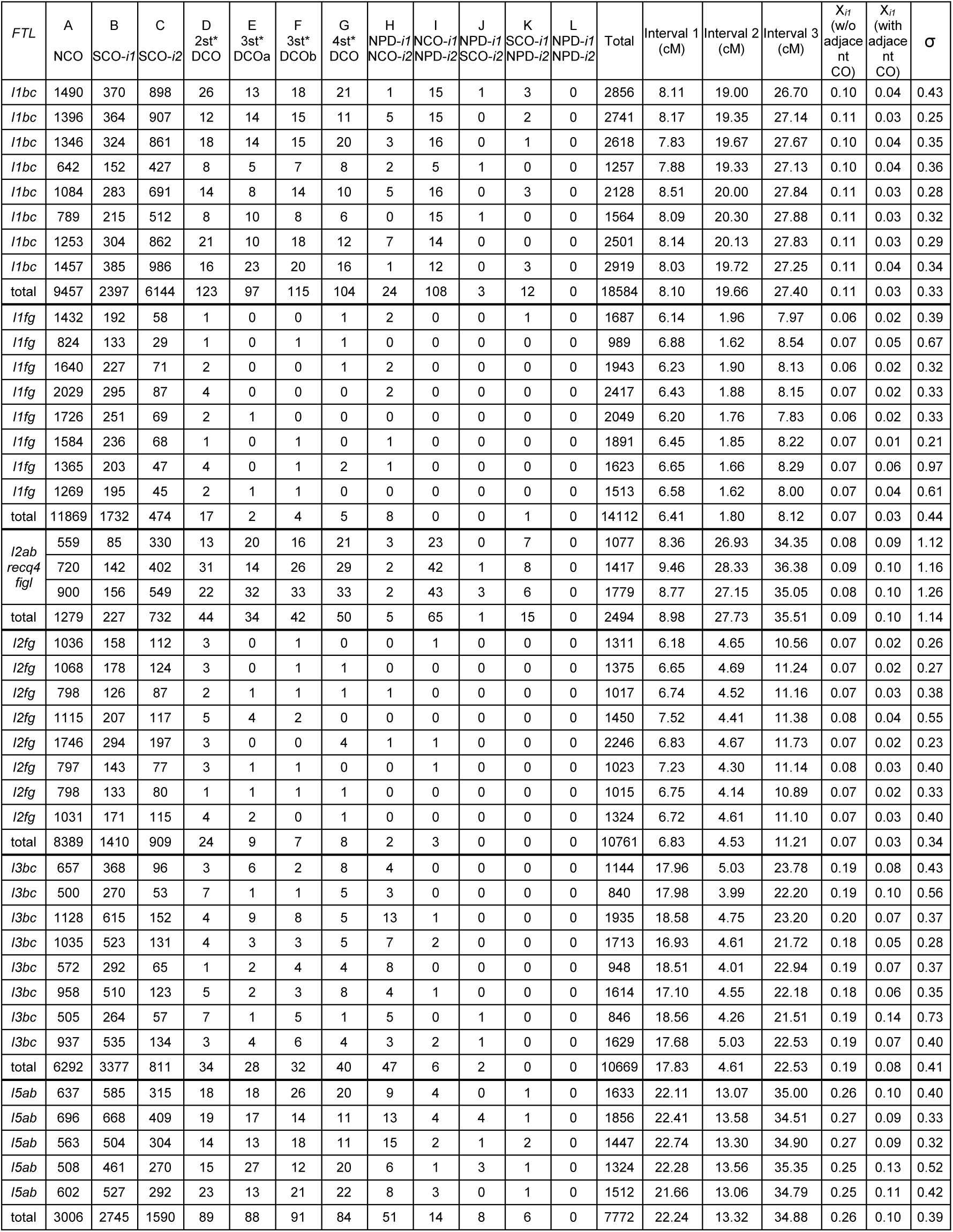
Measurements of crossover frequency and interference in FTL intervals by DeepTetrad. Abbreviations: CO, crossover; non-crossover, NCO; single crossover, SCO; double crossover, DCO; st*, stand; NPD, non-parental ditype; σ, interference ratio. σ= X*_i1_* (with adjacent CO)/ X*_i1_* (w/o adjacent CO). X *_i1_* is the map distance of the first interval (*i1*) generated from the Perkins equation ((1/2*T)+3*(NPD)/total). A σ value of 1 indicates no interference. The letters A-L represent tetrad classification as described previously^18^.

**Supplementary Table 4.** Measurements of crossover frequency in FTL intervals by manually counting tetrads. Abbreviations: NPD, non-parental ditype; T, tetra type; FTL, fluorescence tagged line.

| FTL Interval | NPD | T | Total | cM | FTL Interval | NPD | T | Total | cM |
| --- | --- | --- | --- | --- | --- | --- | --- | --- | --- |
| CEN3 | 1 | 163 | 670 | 12.61 | I2g | 0 | 11 | 153 | 3.59 |
| CEN3 | 0 | 134 | 517 | 12.96 | I2g | 0 | 9 | 114 | 3.95 |
| CEN3 | 0 | 164 | 623 | 13.16 | I2g | 0 | 6 | 77 | 3.90 |
| CEN3 | 2 | 124 | 528 | 12.88 | I2g | 0 | 15 | 169 | 4.44 |
| CEN3 | 2 | 155 | 593 | 14.08 | I2g | 0 | 9 | 119 | 3.78 |
| CEN3 | 0 | 124 | 563 | 11.01 | I2g | 0 | 10 | 99 | 5.05 |
| CEN3 | 2 | 150 | 656 | 12.35 | I2g | 0 | 22 | 205 | 5.37 |
| I1b | 0 | 151 | 998 | 7.57 | I2f-g | 0 | 119 | 547 | 10.88 |
| I1b | 0 | 112 | 764 | 7.33 | I2f-g | 1 | 88 | 363 | 12.95 |
| I1b | 0 | 119 | 737 | 8.07 | I2f-g | 2 | 126 | 556 | 12.41 |
| I1b | 0 | 118 | 692 | 8.53 | I2f-g | 0 | 65 | 333 | 9.76 |
| I1b | 0 | 35 | 224 | 7.81 | I3b | 1 | 190 | 542 | 18.08 |
| I1b | 0 | 118 | 670 | 8.81 | I3b | 2 | 126 | 380 | 18.16 |
| I1c | 3 | 347 | 998 | 18.29 | I3b | 4 | 230 | 694 | 18.30 |
| I1c | 2 | 234 | 618 | 19.90 | I3b | 4 | 64 | 228 | 19.30 |
| I1c | 8 | 261 | 737 | 20.96 | I3b | 9 | 273 | 784 | 20.85 |
| I1c | 4 | 258 | 710 | 19.86 | I3b | 0 | 59 | 193 | 15.28 |
| I1c | 9 | 180 | 577 | 20.28 | I3b | 1 | 68 | 219 | 16.89 |
| I1c | 6 | 182 | 521 | 20.92 | I3b | 2 | 99 | 296 | 18.75 |
| I1c | 4 | 174 | 500 | 19.80 | I3b | 4 | 167 | 559 | 17.08 |
| I1b-c | 6 | 472 | 998 | 25.45 | I3b | 2 | 134 | 381 | 19.16 |
| I1b-c | 4 | 435 | 764 | 30.04 | I3b | 1 | 65 | 207 | 17.15 |
| I1b-c | 10 | 352 | 737 | 27.95 | I3b | 0 | 59 | 184 | 16.03 |
| I1b-c | 4 | 387 | 674 | 30.49 | I3b | 1 | 67 | 196 | 18.62 |
| I1b-c | 2 | 120 | 213 | 30.99 | I3b | 0 | 110 | 311 | 17.68 |
| I1b-c | 2 | 45 | 132 | 21.59 | I3b | 2 | 126 | 391 | 17.65 |
| I1b-c | 0 | 25 | 47 | 26.60 | I3b | 2 | 166 | 494 | 18.02 |
| I1f | 0 | 56 | 536 | 5.22 | I3c | 1 | 22 | 276 | 5.07 |
| I1f | 0 | 56 | 494 | 5.67 | I3c | 0 | 20 | 219 | 4.57 |
| I1f | 0 | 35 | 292 | 5.99 | I3c | 0 | 31 | 296 | 5.24 |
| I1f | 1 | 70 | 589 | 6.45 | I3c | 1 | 50 | 559 | 5.01 |
| I1f | 0 | 61 | 607 | 5.02 | I3c | 0 | 34 | 381 | 4.46 |
| I1f | 18 | 1 | 169 | 7.10 | I3c | 0 | 18 | 184 | 4.89 |
| I1f | 29 | 0 | 222 | 6.53 | I3c | 0 | 21 | 196 | 5.36 |
| I1f | 23 | 0 | 198 | 5.81 | I3c | 0 | 29 | 311 | 4.66 |
| I1f | 0 | 16 | 129 | 6.20 | I3c | 0 | 33 | 391 | 4.22 |
| I1f | 0 | 22 | 199 | 5.53 | I3c | 0 | 53 | 494 | 5.36 |
| I1f | 0 | 17 | 152 | 5.59 | I3b-c | 5 | 92 | 233 | 26.18 |
| I1f | 0 | 18 | 188 | 4.79 | I3b-c | 3 | 91 | 219 | 24.89 |
| I1f | 0 | 21 | 196 | 5.36 | I3b-c | 4 | 122 | 296 | 24.66 |
| I1f | 0 | 22 | 155 | 7.10 | I3b-c | 6 | 208 | 559 | 21.82 |
| I1f | 0 | 12 | 99 | 6.06 | I3b-c | 6 | 162 | 381 | 25.98 |
| I1f | 0 | 15 | 122 | 6.15 | I3b-c | 0 | 74 | 184 | 20.11 |
| I1f | 0 | 7 | 118 | 2.97 | I3b-c | 2 | 85 | 196 | 24.74 |
| I1g | 0 | 17 | 645 | 1.32 | I3b-c | 2 | 135 | 311 | 23.63 |
| I1g | 0 | 22 | 532 | 2.07 | I3b-c | 4 | 143 | 391 | 21.36 |
| I1g | 0 | 17 | 294 | 2.89 | I3b-c | 4 | 200 | 494 | 22.67 |
| I1g | 1 | 36 | 739 | 2.84 | I5a | 2 | 266 | 608 | 22.86 |
| I1g | 0 | 31 | 665 | 2.33 | I5a | 6 | 208 | 513 | 23.78 |
| I1g | 0 | 21 | 500 | 2.10 | I5a | 2 | 48 | 139 | 21.58 |
| I1f-g | 0 | 70 | 544 | 6.43 | I5a | 1 | 34 | 99 | 20.20 |
| I1f-g | 0 | 74 | 475 | 7.79 | I5a | 1 | 53 | 140 | 21.07 |
| I1f-g | 0 | 43 | 256 | 8.40 | I5b | 0 | 168 | 608 | 13.82 |
| I2f | 0 | 68 | 532 | 6.39 | I5b | 0 | 102 | 407 | 12.53 |
| I2f | 0 | 53 | 363 | 7.30 | I5b | 0 | 119 | 442 | 13.46 |
| I2f | 1 | 82 | 556 | 7.91 | I5b | 0 | 107 | 493 | 10.85 |
| I2f | 0 | 41 | 333 | 6.16 | I5b | 1 | 30 | 136 | 13.24 |
| I2f | 0 | 91 | 525 | 8.67 | I5b | 0 | 20 | 99 | 10.10 |
| I2g | 0 | 47 | 480 | 4.90 | I5b | 1 | 29 | 140 | 12.50 |
| I2g | 0 | 46 | 363 | 6.34 | I5a-b | 9 | 369 | 608 | 34.79 |
| I2g | 0 | 50 | 556 | 4.50 | I5a-b | 18 | 304 | 605 | 34.05 |
| I2g | 0 | 23 | 334 | 3.44 | I5a-b | 11 | 281 | 476 | 36.45 |
| I2g | 0 | 59 | 536 | 5.50 | I5a-b | 8 | 280 | 493 | 33.27 |
| I2g | 0 | 71 | 639 | 5.56 | I5a-b | 2 | 90 | 134 | 38.06 |
| I2g | 0 | 12 | 126 | 4.76 | I5a-b | 1 | 59 | 99 | 32.83 |
| I2g | 0 | 6 | 73 | 4.11 | I5a-b | 1 | 78 | 140 | 30.00 |

**Supplementary Table 5.** Measurements of crossover interference in FTL-*I1bc* by manually counting tetrads. Abbreviations: CO, crossover; non-crossover, NCO; single crossover, SCO; double crossover, DCO; st*, stand; NPD, non-parental ditype; σ, interference ratio. σ= X*_i1_* (with adjacent CO)/ X*_i1_* (w/o adjacent CO). X *_i1_* is the map distance of the first interval (*i1*) generated from the Perkins equation ((1/2*T)+3*(NPD)/total). A σ value of 1 indicates no interference. The letters A-L represent tetrad classification as described previously^18^.

| FTL | A<br>NCO | B<br>SCO-*i1* | C<br>SCO-*i2* | D<br>2st*<br>DCO | E<br>3st*<br>DCOa | F<br>3st*<br>DCOb | G<br>4st*<br>DCO | H<br>NPD-*i1*<br>NCO-*i2* | I<br>NCO-*i1*<br>NPD-*i2* | J<br>NPD-*i1*<br>SCO-*i2* | K<br>SCO-*i1*<br>NPD-*i2* | L<br>NPD-*i1*<br>NPD-*i2* | Total | Interval 1<br>(cM) | Interval 2<br>(cM) | Interval 3<br>(cM) | $X_{i1}$ (w/o<br>adjacent<br>CO) | $X_{i1}$ (with<br>adjacent<br>CO) | $\sigma$ |
| --- | --- | --- | --- | --- | --- | --- | --- | --- | --- | --- | --- | --- | --- | --- | --- | --- | --- | --- | --- |
| *l1bc* | 514 | 110 | 277 | 7 | 4 | 4 | 4 | 0 | 3 | 0 | 0 | 0 | 923 | 6.99 | 17.01 | 23.67 | 0.09 | 0.03 | 0.36 |

**Supplementary Table 6.** Information of imaging fluorescent pollen in *FTLs.* The gain, saturation, gamma values except for the exposure value are applied across images: gain of x10.1, saturation of 1.00, gamma of 1.50. In particular, exposure for the YFP image should be adjusted if its fluorescence is weaker than those in the other channels. The table shows the value of exposure applied to each FTL (fluorescent tagged line).

| FTL | Red<br>(ms) | Yellow<br>(ms) | Cyan<br>(ms) |
| --- | --- | --- | --- |
| *I1bc (YRC/+++)* | 250 | 1,200 | 1,200 |
| *I1fg (YCR/+++)* | 300 | 800 | 800 |
| *I2ab (CYR/+++)* | 150 | 700 | 1,500 |
| *I2fg (RYC/+++)* | 40 | 1500 | 700 |
| *I3bc(CYR/+++)* | 300 | 1300 | 800 |
| *CEN3 (YR/++)* | 700 | 1500 |  |
| *I5ab (RYC/+++)* | 500 | 1,500 | 1500 |

## References

1. Hunter, N. Meiotic Recombination: The Essence of Heredity. Cold Spring Harb. Perspect. Biol. 7, a016618 (2015).

2. Mercier, R., Mézard, C., Jenczewski, E., Macaisne, N. & Grelon, M. The Molecular Biology of Meiosis in Plants. Annu. Rev. Plant Biol. 66, 297–327 (2015).

3. Wang, Y. & Copenhaver, G. P. Meiotic Recombination: Mixing It Up in Plants. Annu. Rev. Plant Biol. 69, 577–609 (2018).

4. Vrielynck, N. et al. A DNA topoisomerase VI-like complex initiates meiotic recombination. Science 351, 939–43 (2016).

5. Choi, K. & Henderson, I. R. Meiotic recombination hotspots - A comparative view. Plant J. 83, (2015).

6. Choi, K. et al. Nucleosomes and DNA methylation shape meiotic DSB frequency in Arabidopsis thaliana transposons and gene regulatory regions. Genome Res. 28, (2018).

7. Choi, K. et al. Arabidopsis meiotic crossover hot spots overlap with H2A.Z nucleosomes at gene promoters. Nat. Genet. 45, (2013).

8. Mercier, R. et al. Two meiotic crossover classes cohabit in Arabidopsis: one is dependent on MER3,whereas the other one is not. Curr. Biol. 15, 692–701 (2005).

9. Higgins, J. D., Armstrong, S. J., Franklin, F. C. H. & Jones, G. H. The Arabidopsis MutS homolog AtMSH4 functions at an early step in recombination: evidence for two classes of recombination in Arabidopsis. Genes Dev. 18, 2557–70 (2004).

10. Berchowitz, L. E., Francis, K. E., Bey, A. L. & Copenhaver, G. P. The role of AtMUS81 in interference-insensitive crossovers in A. thaliana. PLoS Genet. 3, e132 (2007).

11. Serra, H. et al. Massive crossover elevation via combination of HEI10 and recq4a recq4b during Arabidopsis meiosis. Proc. Natl. Acad. Sci. U. S. A. 115, 2437–2442 (2018).

12. Fernandes, J. B., Séguéla-Arnaud, M., Larchevêque, C., Lloyd, A. H. & Mercier, R. Unleashing meiotic crossovers in hybrid plants. Proc. Natl. Acad. Sci. U. S. A. 115, 2431–2436 (2018).

13. Choi, K. Advances towards Controlling Meiotic Recombination for Plant Breeding. Mol. Cells 40, (2017).

14. Crismani, W. et al. FANCM limits meiotic crossovers. Science 336, 1588–90 (2012).

15. Girard, C. et al. AAA-ATPase FIDGETIN-LIKE 1 and Helicase FANCM Antagonize Meiotic Crossovers by Distinct Mechanisms. PLoS Genet. 11, e1005369 (2015).

16. Mieulet, D. et al. Unleashing meiotic crossovers in crops. Nat. Plants 4, 1010–1016 (2018).

17. Francis, K. E. et al. Pollen tetrad-based visual assay for meiotic recombination in Arabidopsis. Proc. Natl. Acad. Sci. U. S. A. 104, 3913–8 (2007).

18. Berchowitz, L. E. & Copenhaver, G. P. Fluorescent Arabidopsis tetrads: a visual assay for quickly developing large crossover and crossover interference data sets. Nat. Protoc. 3, 41–50 (2008).

19. Yelina, N. E. et al. High-throughput analysis of meiotic crossover frequency and interference via flow cytometry of fluorescent pollen in Arabidopsis thaliana. Nat. Protoc. 8, 2119–2134 (2013).

20. Ziolkowski, P. A. et al. Juxtaposition of heterozygosity and homozygosity during meiosis causes reciprocal crossover remodeling via interference. Elife 4, e03708 (2015).

21. Ziolkowski, P. A. et al. Natural variation and dosage of the HEI10 meiotic E3 ligase control Arabidopsis crossover recombination. Genes Dev. 31, 306–317 (2017).

22. He, K., Zhang, X., Ren, S. & Sun, J. Deep Residual Learning for Image Recognition. ArXiv 1512.03385 (2015).

23. He, K., Gkioxari, G., Dollár, P. & Girshick, R. Mask R-CNN. ArXiv 1703.06870 (2017).

24. Lin, T.-Y. et al. Feature Pyramid Networks for Object Detection. ArXiv1612.03144 (2016).

25. Abadi, M. et al. TensorFlow: A system for large-scale machine learning. in Operating Systems Design and Implementation 265–283 (2016).

26. Chollet, F. keras. GitHub (2015).

27. Giraut, L. et al. Genome-wide crossover distribution in Arabidopsis thaliana meiosis reveals sex-specific patterns along chromosomes. PLoS Genet. 7, (2011).

28. Girshick, R. Fast R-CNN. ArXiv 1504.08083 (2015).

29. Lin, T.-Y. et al. Microsoft COCO: Common Objects in Context. ArXiv 1405.0312v3 (2015).

30. Dutta, A. & Zisserman, A. The VGG Image Annotator (VIA). ArXiv 1904.10699v1 (2019).

31. Yelina, N. E. et al. DNA methylation epigenetically silences crossover hot spots and controls chromosomal domains of meiotic recombination in Arabidopsis. Genes Dev. 29, 2183–202 (2015).

